# Endothelial PHD2 Deficiency Induces Nitrative Stress via Suppression of Caveolin-1 in Pulmonary Arterial Hypertension

**DOI:** 10.1101/2021.09.24.461744

**Authors:** Bin Liu, Yi Peng, Dan Yi, Jingbo Dai, Rebecca Vanderpool, Maggie M. Zhu, Zhiyu Dai, You-Yang Zhao

## Abstract

Nitrative stress is a characteristic feature of the pathology of human pulmonary arterial hypertension (PAH). However, the role of nitrative stress in the pathogenesis of obliterative vascular remolding and severe PAH remains largely unclear. Our recent studies identified a novel mouse model (*Egln1*^*Tie2Cre*^, *Egln1* encoding prolyl hydroxylase 2 [PHD2]) with obliterative vascular remodeling and right heart failure, which provides us an excellent model to study the role of nitrative stress in obliterative vascular remodeling. Here we show that nitrative stress was markedly elevated whereas endothelial Caveolin-1 (Cav1) expression was suppressed in the lungs of *Egln1*^*Tie2Cre*^ mice. Treatment with a superoxide dismutase mimetic, manganese (III) tetrakis (1-methyl-4-pyridyl) porphyrin pentachloride (MnTmPyP, also known as a peroxynitrite scavenger) treatment inhibited obliterative pulmonary vascular remodeling and attenuated severe PAH in *Egln1*^*Tie2Cre*^ mice. Genetic restoration of endothelial Cav1 expression in *Egln1*^*Tie2Cre*^ mice normalized nitrative stress, reduced PAH and improved right heart function. These data suggest that suppression of endothelial Cav1 expression secondary to PHD2 deficiency augments nitrative stress, which contributes to obliterative vascular remodeling and severe PAH. Thus, reactive oxygen/nitrogen species scavenger might have great therapeutic potential for the inhibition of obliterative vascular remodeling and severe PAH.

## Introduction

Pulmonary arterial hypertension (PAH) is a devastating disease characterized by persistent increase of pulmonary vascular resistance and obstructive pulmonary vascular remodeling, which lead to right-sided heart failure and premature death (1–4). Histopathological features of PAH from different etiologies include increased vessel wall thickness, vascular fibrosis, augmented oxidative/nitrative stress, microvascular occlusion and complex plexiform lesions (5–7). Given the molecular mechanisms of obliterative pulmonary vascular remodeling are not fully understood, the current PAH therapies targeting mainly vasoconstriction with little effect on vascular remodeling are not effective in curing the disease and promoting survival (8, 9).

There are accumulating evidence of oxidative/nitrative stress in the lungs of IPAH patients. Xanthine oxidase activity involved in the generation of superoxide anions (reactive oxygen species, ROS) is markedly increased in IPAH lungs(10), and they contributes to chronic hypoxia-induced PH(11). NADPH oxidases including Nox1, 2, and 4, which also produce high levels of ROS, have been demonstrated to be involved in the development of PAH(12). 8-Hydroxyguanosine (a biomarker of oxidative damage caused by reaction of superoxide with guanine) staining is prominent in endothelial cells (ECs) within the plexiform lesions of IPAH patients(13). In contrary, antioxidants such as manganese superoxide dismutase (MnSOD, a key mitochondrial antioxidant enzyme coded by the SOD2 gene) activity was lower in IPAH lungs (13, 14). Excessive production of nitric oxide (NO) and superoxide leads to formation of the damaging reactive nitrogen species (RNS) peroxynitrite (15, 16). Elevated levels of peroxynitrite (i.e., nitrative stress) are detrimental, which induce cell damage and death of pulmonary vascular cells, including ECs and smooth muscle cells(12, 17, 18). In addition, peroxynitrite modifies amino acids resulting in tyrosine nitration of proteins (formation of 3-nitrotyrosine) and thus modifies their functions (15, 16, 19). Peroxynitrite levels are elevated in the lungs of PAH patients (13, 14), which are in part due to tissue hypoxia and inflammation (12, 17). Immunohistochemical staining show ubiquitous and marked increase of 3-nitrotyrosine (13). Our previous studies showed that Caveolin-1 (Cav1) deficiency results in eNOS activation and a marked increase of nitric oxide levels, as well as augmented production of superoxide, which form peroxynitrite leading to prominent nitrative stress in pulmonary vasculature (20–22). The resultant augmentation of nitrative stress causes protein kinase G (PKG) tyrosine nitration and impairs its kinase activity, leading to enhanced vasoconstriction and vascular remodeling and thus PH as seen in *Prkg*^*-/-*^(*Prkg* encodes PKG) mice (21–23). PKG tyrosine nitration was markedly increased in the lungs of IPAH patients (21, 24).

Recently, we have reported the unprecedented mouse model of PAH [*Tie2Cre*-mediated disruption of *Egln1*, encoding hypoxia inducible factor (HIF) prolyl hydroxylase 2 (PHD2), designated *Egln1*^*Tie2Cre*^, CKO] with progressive obliterative vascular remodeling including vascular occlusion and plexiform-like lesion and right heart failure, which recapitulates many features of clinical PAH including IPAH(25, 26). It is unknown whether oxidative/nitrative stress is also augmented and involved in the pathogenesis of PAH in this severe PAH model. In the present study, we for the first time demonstrate endothelial PHD2 deficiency induces extensive oxidative/nitrative stress via downregulation of endothelial Cav1 expression in a HIF-2α dependent manner. ROS scavenger treatment or restoration of endothelial Cav1 expression attenuated PAH in CKO mice.

## Methods

### Animal Models

All the mice used in this study are C57BL/6J background. *Egln1*^*Tie2Cre*^ (CKO) and *Hif2a*/*Egln1*^*Tie2Cre*^ (EH2) mice were generated as described previously(25). *Cav1*^EC-Tg^ mice were obtained from Dr. William Sessa’s lab from Yale University, *Cav1*^EC-Tg^ mice were generated by using SpeI-linearized 9.2-kb prepro-endothelin-1 promoter to drive the Cav-1 transgene.(27, 28). CKO mice were bred with Cav1^EC-Tg^ mice to generate *Egln1*^*Tie2Cre*^/*Cav1*^EC-Tg^ (CKO/Tg) and *Egln1*^*f/f*^/*Cav1*^EC-Tg^ (Tg) mice. Both male and female *Egln1*^*f/f*^ (designed as WT), CKO, CKO/Tg, and Tg little mates at the age of 5 weeks to 3.5 months were used in these studies. For ROS scavenger treatment, both male and female CKO mice at the age of 5 weeks were treated with manganese (III) tetrakis (1-methyl-4-pyridyl) porphyrin pentachloride (MnTmPyP, Mn) (5 mg/kg) (21) intraperitoneally daily for 9 weeks. PBS was used as a control. The same PBS group was used in our previous publication (29). The animal care and study protocols were reviewed and approved by the Institutional Animal Care and Use Committees of the University of Illinois at Chicago, Northwestern University, and the University of Arizona.

### Echocardiography

Echocardiography was performed in the Core facility at the University of Illinois at Chicago, Northwestern University Feinberg School of Medicine and the University of Arizona as described previously(25, 30, 31). Transthoracic echocardiography was performed on a VisualSonics Vevo 2100 or 3100 ultrasound machine (FujiFilm VisualSonics Inc) using an MS550D (40 MHz) transducer. The right ventricle wall thickness during diastole (RVWTD) was obtained from the parasternal short axis view at the papillary muscle level using M-mode. The RV cross-sectional area was obtained from the parasternal short axis view at the papillary muscle level using B-mode. Pulmonary artery (PA) acceleration time and PA ejection time were obtained from the parasternal short axis view at the aortic valve level using pulsed Doppler mode. The left ventricle fractional shortening (LV FS) and the cardiac output (CO) were obtained from the parasternal short axis view using M-mode.

### Hemodynamics measurement

In order to measure the right ventricular systolic pressure (RVSP), mice were anesthetized with Ketamine/Xylazine cocktail. A 1.4F pressure transducer catheter (Millar Instruments) was inserted through the right jugular vein into the right ventricle. RVSP was recorded and analyzed by AcqKnowledge software (Biopac Systems Inc.) as described previously(25, 30, 31).

### Endothelial cells isolation

Mouse lung tissues were well-perfused with PBS to remove blood. The tissues were then minced and incubated with collagenase A (1.0 mg/ml in HBSS, Roche) in water bath shaker at 37°C for 45 mins, and then were dispersed by gentleMACS Dissociator (Miltenyi Biotec), followed by filtering through a 40 μm cell strainer. Cells were then incubated with anti-mouse CD31 antibody (BD, cat#550274, 1.5 μg/ml) on ice for 1 hour and followed by incubation with sheep anti-rat IgG Dynabeads M-450 (Invitrogen, Cat#11035, 50 μl) for 30 mins. Beads bound ECs (CD31^+^ cells) were pulled down by magnet (27, 28) and lysed for protein isolation.

### Histological assessment

Mouse lung tissues were perfused with PBS and fixed with 10% formalin via tracheal instillation at a constant pressure (15 cm H_2_O) and embedded in paraffin wax. Lung sections were stained with a Russel-Movat pentachrome staining kit (American MasterTech) according to the manufacturer’s protocols. For assessment of PA wall thickness, PAs from 40 images at 20X magnification were quantified by Image J. Wall thickness was calculated by the distance between internal wall and external wall divided by the distance between external wall and the center of lumen (24, 27, 28).

### Immunofluorescent staining

Mouse lung tissues were perfused with PBS, inflated with 50% OCT in PBS, and embedded in OCT for cryosectioning. For immunofluorescent staining of Cav1 and α-nitrotyrosine (NT) (20), lung sections (5 μm) were fixed with 4% paraformaldehyde and blocked with 0.1% Triton X-100 and 5% normal goat serum at room temperature for 1 hour. After 3 washes with PBS, the slides were incubated with anti-Cav1 (Santa Cruz Biotechnology, Cat#sc-894, 1:100) or anti-NT (Millipore, Cat#05-233,1:300), anti-CD31 antibody (BD Bioscience, Cat#550274, 1:25), anti-α-SMA (Abcam, Cat#Ab5694, 1:300), anti-CD45 (Invitrogen, Cat# 14-0451-82, 1:100), anti-Periostin (Abcam, Cat#14041, 1:100) at 4°C overnight then incubated with Alexa 594-conjugated anti-mouse IgG, Alexa 488 or 647-conjugated anti-rat or anti-rabbit IgG (Life Technology) at room temperature for 1 h. Nuclei were counterstained with DAPI contained in Prolong Gold mounting media (Life Technology).

### RNA isolation and quantitative RT-PCR analysis

One lobe of lung tissue was homogenized by a TissueLyser (Qiagen) in Trizol solution (Life Technology). RNA was isolated by phenol:chloroform, followed by cleanup with Qiagen RNA mini kit (Qiagen). One microgram of RNA was transcribed into cDNA using the high-capacity cDNA reverse transcription kits (Applied Biosystems). Quantitative RT-PCR analysis was performed on an ABI ViiA 7 Real-time PCR system (Applied Biosystems) with the FastStart SYBR Green Master kit (Roche Applied Science). Target mRNA expression was determined using the comparative cycle threshold method of relative quantitation. Cyclophilin A was used as an internal control. Information for mouse Cav1 and Cyclophilin A primers was described previously(25).

### Immunoprecipitation and Western Blotting

Lung tissues were collected from 3.5 months old mice for homogenization. 500 μg of lysates/mouse were then immunoprecipitated with anti-PKG antibody(19) and protein conjugated-Sepharose beads and blotted with anti-NT antibody (Millipore, Cat#05-233, 1:2000). The same membrane was also blotted with anti-PKG antibody as a loading control. Other Western blotting assays were performed using anti-Cav1 (Santa Cruz Biotechnology, Cat#sc-894 or sc-53564, 1:2000), anti-NT (Millipore, Cat#05-233, 1:2000). Anti-β-actin (Sigma, Cat #A2228, 1:10,000) was used as a loading control.

### Measurement of reactive oxygen/nitrogen species (ROS/RNS) levels

Lung tissues were perfused free of blood with PBS, followed by collection of left lobe. Fifty mg of lung tissue was homogenized in PBS on ice. Tissue homogenates were span at 10,000g for 5 min in 4°C. Fifty ul of homogenates were then used for determination of tissue ROS/RNS levels (OxiSelect™ In Vitro ROS/RNS Assay Kit, Cat#STA-347-T) according to the manufacture’s guide. 2’, 7’-dichlorodihydrofluorescein (DCF) in the samples were measured fluorometrically against DCF standards on a fluorescence plate reader (TECAN Infinite 200 Pro). The free radical content in homogenates is determined based on DCF standard curve. The relative DCF values were normalized by the mean value of WT group.

### Statistical analysis

Statistical significance was determined by one-way ANOVA with a Tukey post hoc analysis that calculates P values corrected for multiple comparisons. Two-group comparisons were analyzed by the unpaired 2-tailed Student t-test. P <0.05 denoted the presence of a statistically significant difference. All bar graphs represent mean ± SD.

## Results

### Prominent nitrative stress in *Egln1*^*Tie2Cre*^ mouse lungs and pulmonary vascular ECs

Given that nitrative stress is a characteristic feature of the pathology of clinical PAH, we tested whether nitrative stress is also prominent in the lung of *Egln1*^*Tie2Cre*^ (CKO) mice. Quantification of total free radical levels with the OxiSelect™ *In Vitro* ROS/RNS Assay Kit of lung homogenates showed that CKO lungs exhibited markedly elevated ROS/RNS levels (**Figure 1A**). Western blotting demonstrated that nitrotyrosine (NT) expression, indicative of nitrative stress (12), was strongly induced in CKO mouse lungs (**Figure 1B** and **1C**). We then performed the NT staining together with markers for ECs (CD31), inflammatory cells (CD45), fibroblasts (periostin), smooth muscle cells (α-SMA) to determine what cell types are the NT positive cells in the lung of CKO mice. Our data showed that NT was highly expressed in the pulmonary vascular ECs, CD45^+^ inflammatory cells and relatively low expression in pulmonary vascular smooth muscle cells and periostin^+^ fibroblasts (**Figure 1D-1E and supplemental Figure 1A and 1B**). Our previous studies demonstrated that nitration–dependent impairment of PKG activity is involved in PH development(19). To further determine whether PHD2 deficiency also affects PKG nitration and expression, we performed immunoprecipitation assay using anti-PKG antibodies and blotted with anti-NT antibodies. The data showed that PKG nitration was markedly increased in the lung of CKO mice (**Supplemental Figure 1C**). Together, these data demonstrate that nitrative stress is markedly augmented in the pulmonary vascular lesions in CKO mice.

**Figure 1.**
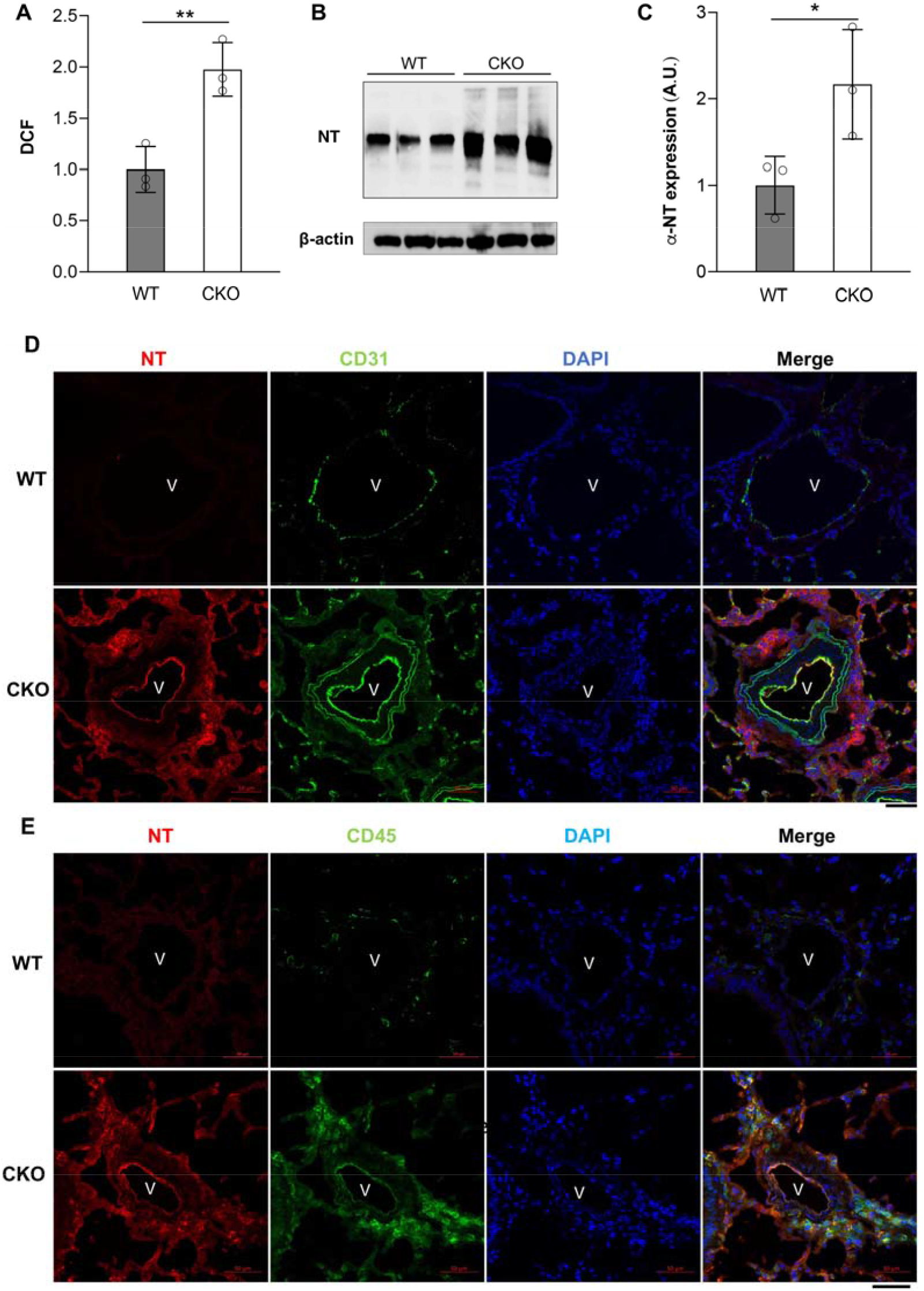
Augmented nitrative stress in pulmonary vascular lesions in *Egln1*^*Tie2Cre*^ mice. (**A**) DCF measurement demonstrating marked increase of ROS/RNS in the lungs of *Egln1*^*Tie2Cre*^ (CKO) mice. Lung tissues were collected from 3.5 months old WT and CKO mice for 2’, 7’-dichlorodihydrofluorescein (DCF) assay. (**B, C**) Western blotting demonstrating increase of nitrotyrosine modification of proteins in the lung of CKO mice. NT, anti-NT antibody. Anti-β-actin was used as a control. (**D, E**) Representative micrographs of immunostaining showing prominent nitrative stress in pulmonary vascular lesions in CKO mice (3.5 months old). Lung sections were immunostained with anti-NT antibody (red) for detection of nitrative stress. Anti-CD31 antibodies were used to label vascular ECs (green, **D**). Anti-CD45 antibodies were used to label inflammatory cells (green, **E**). Nuclei were counterstained with DAPI (blue). V, vessel. Scale bar, 50 μm. *, *P*< 0.05. **, *P* < 0.01. Student t-test. (**A, C**).

### ROS scavenger treatment inhibited obliterative vascular remodeling and attenuate PAH in *Egln1*^*Tie2*Cre^ mice

We next determined whether removing ROS/RNS in CKO mice will affect the development of PAH. CKO mice at age of 5-weeks (with established PAH phenotype) were treated with the ROS/peroxynitrite scavenger MnTMPyP (Mn), which disrupts superoxide and thus peroxynitrite formation (21, 32), via i.p. injection for 9 weeks (**Figure 2A**). Mn treatment markedly reduced the levels of NT in the pulmonary vasculature and lung of CKO mice (**Figure 2B**). Hemodynamics measurement showed that RVSP was reduced to ∼50 mmHg in Mn-treated CKO mice compared to ∼77 mmHg in PBS-treated CKO mice (**Figure 2B**). RV hypertrophy, determined by weight ratio of RV versus left ventricular plus septum (LV+S), was also attenuated in Mn-treated CKO mice. Lung pathology examination also revealed inhibition of occlusive vascular remodeling evident by diminished neointima formation (**Figure 3A**), and decreased pulmonary arterial wall thickness and muscularization of distal pulmonary vessels in Mn-treated CKO mice compared to PBS treated mice (**Figure 3B-3E**). Together, these data suggest the involvement of oxidative/nitrative stress in severe PAH in CKO mice.

**Figure 2.**
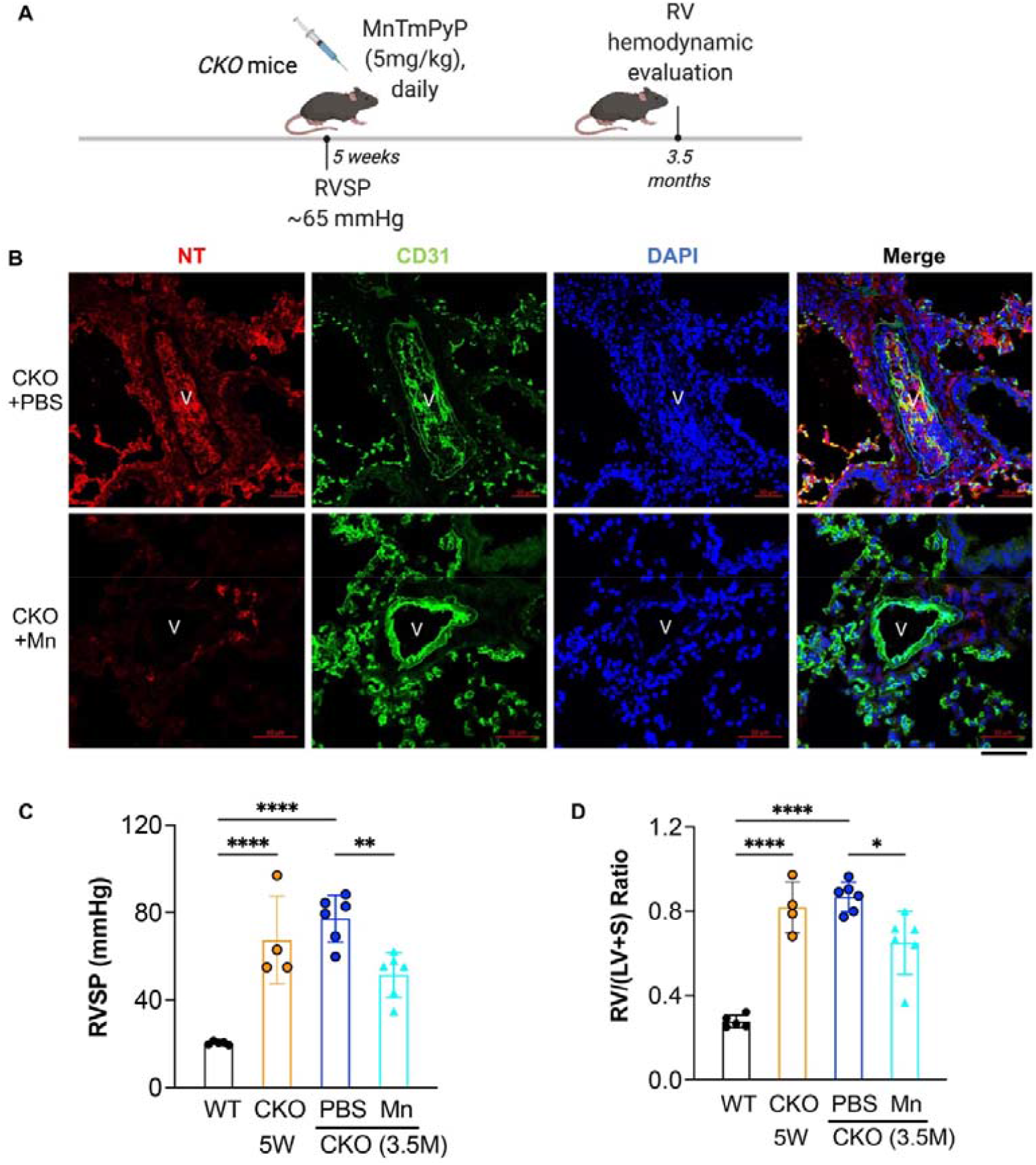
MnTMPyP treatment reduced PAH in *Egln1*^*Tie2Cre*^ mice. (**A**) A diagram showing treatment strategy in *Egln1*^*Tie2Cre*^ (CKO) mice with MnTMPyP (Mn). (B) Immunostaining against NT showed that MnTMPyP (Mn) treatment reduced nitrotyrosine modification in CKO lung. CKO mice at age of 5 weeks were treated with either Mn or PBS for 9 weeks. (**C, D**) RVSP was measured and RV hypertrophy was determined in CKO mice (3.5 months, 3.5M) treated with Mn or PBS. *Egln1*^*f/f*^ (WT) mice were used as controls. RVSP and RV hypertrophy were also determined in 5 weeks (5W) old *CKO* mice without treatment. **, *P* < 0.01, ****, *P* < 0.0001, One-way ANOVA with Tukey post-hoc analysis (**C** and **D**).

**Figure 3.**
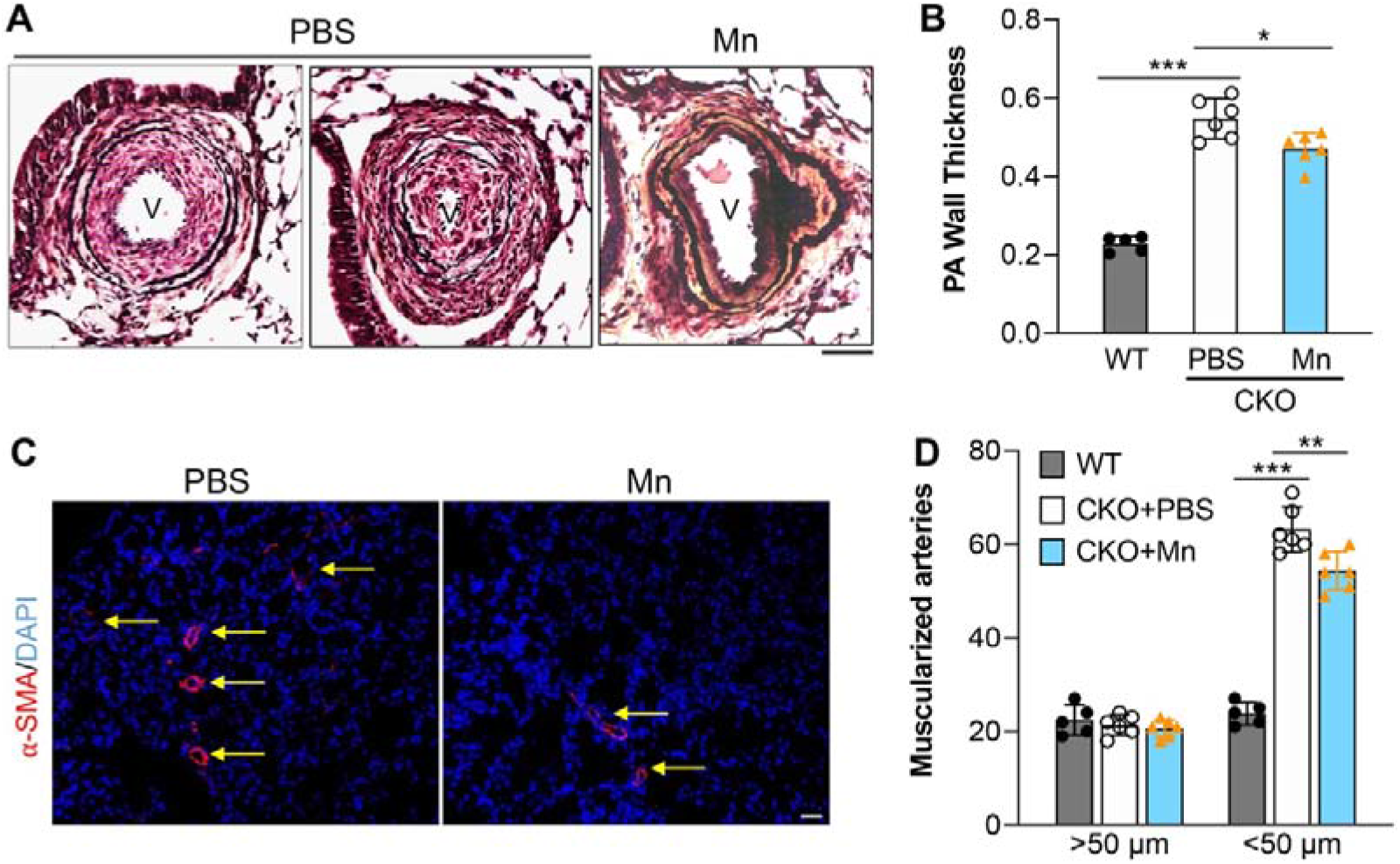
MnTMPyP inhibited obliterative vascular remodeling in *Egln1*^*Tie2Cre*^ mice. (**A, B**) MnTMPyP (Mn) treatment inhibited neointima formation and decreased PA wall thickness. Lung sections were processed for Russel-Movat pentachrome staining. V, vessel. (**C, D**) Quantification of α-SMA immunostaining demonstrating decreased muscularization of distal pulmonary vessels. Lung cryosections were immunostained with anti-α-SMA (red) and counterstained with DAPI (blue). Arrows point to muscularized distal pulmonary vessels. *, *P* < 0.05, **, *P* < 0.01, ***, *P* <0.001. One-way ANOVA with Tukey post-hoc analysis (**B** and **D**). Scale bar: 50μm.

### Decreased expression of endothelial Caveolin-1 secondary to PHD2 deficiency in a HIF-2α-dependent manner

Cav1 deficiency increases ROS and RNS production, leading to nitrative stress, which post-translationally modifies PKG through tyrosine nitration and impairs its kinase activity leading to PH(21, 22). To find out whether dysregulation of Cav1 signaling is involved in the generation of nitrative stress and PAH development in CKO mice, we first checked the expression level of Cav1 in the lungs of CKO mice. As shown in **Figure 4A**, *Cav1* mRNA levels was markedly decreased in the lungs of CKO mice compared to that from similar age *Egln1*^*f/f*^ (WT) mice. HIF-2α deletion in CKO mice restored the expression of *Cav1* mRNA (**Figure 4A**). Western blotting also demonstrated reduced Cav1 protein levels in CKO mouse lungs which were reversed in *Egln1*^*Tie2Cre*^*/Hif2a*^*Tie2Cre*^ (EH2) mice and in fact increased in EH2 mouse lungs compared to WT lungs (**Figure 4B** and **4C**). By performing immunofluorescent staining, we found that Cav1 is co-stained with CD31^+^ cells (i.e. ECs) in the pulmonary vasculature of WT mice whereas diminished in pulmonary vascular lesions in CKO mice and restored in CD31^+^ cells in the pulmonary vascular ECs of EH2 mice (**Figure 4D**), suggesting that endothelial Cav1 is reduced by PHD2 deficiency dependent on HIF-2α activation in *Egln1*^*Tie2Cre*^ mice.

**Figure 4.**
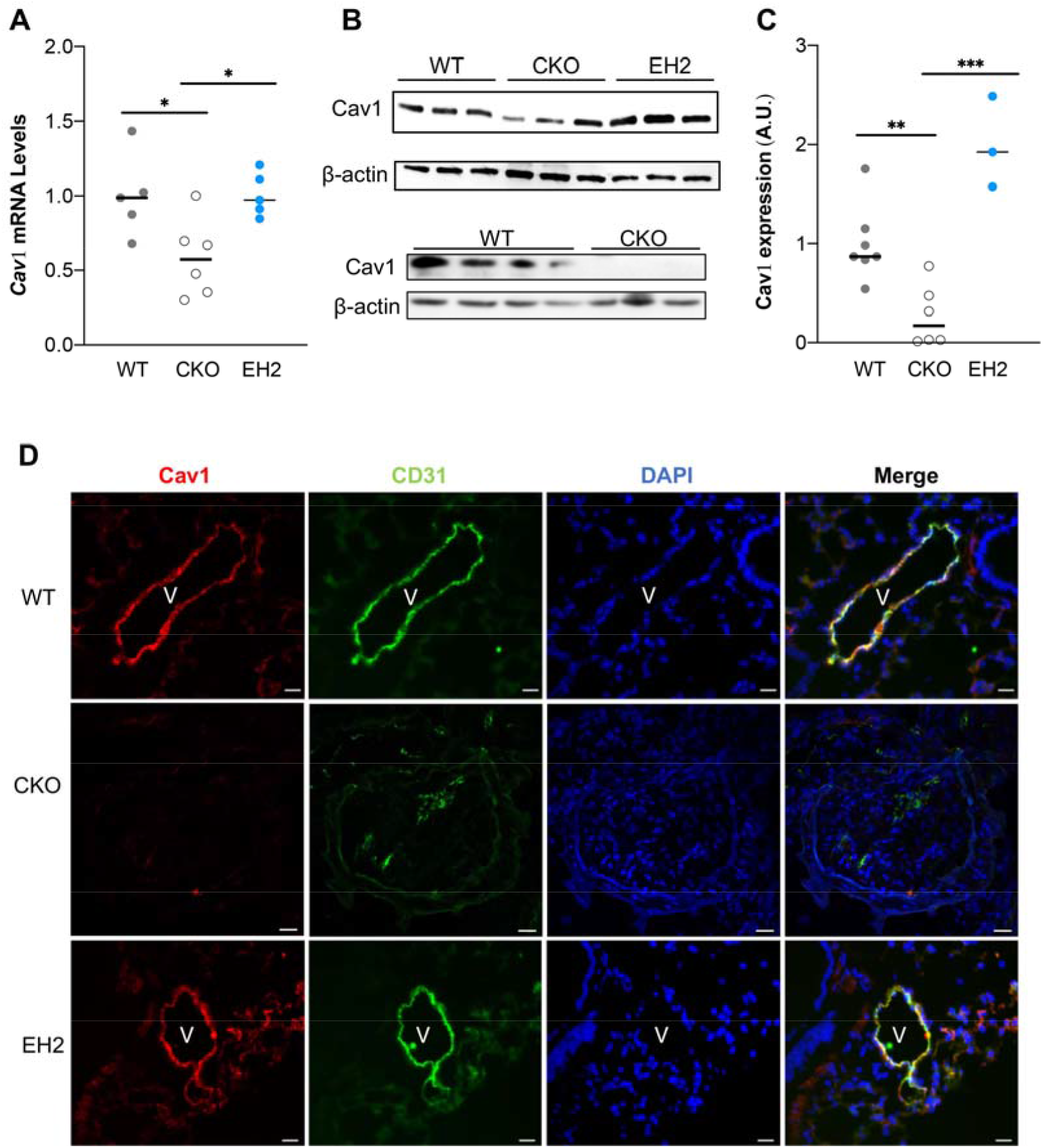
Endothelial Cav1 expression was suppressed in *Egln1*^*Tie2Cre*^ mouse lungs in a HIF-2α dependent manner. (**A**) Quantitative RT-PCR analysis demonstrating decreased Cav1 mRNA expression in *Egln1*^*Tie2Cre*^ (CKO) mouse lungs compared to WT lungs, which is restored in *Egln1*^*Tie2Cre*^/*Hif2a*^*Tie2Cre*^ (EH2) lungs. Lung tissues were collected from 3.5 months old mice. (**B, C**) Decreased Cav1 protein expression in CKO mouse lungs compared to WT lungs. Cav1 protein expression in EH2 mouse lungs was markedly increased compared to WT and CKO mouse lungs. (**D**) Immunostaining demonstrating diminished Cav1 expression in pulmonary vascular ECs of CKO mice and restoration in EH2 mice. Lung sections were immunostained with anti-Cav1 (red) and CD31 (green). Nuclei were counterstained with DAPI (blue). *, *P* < 0.05, **, *P* <0.01, ***, *P* < 0.001. One-way ANOVA with Tukey post-hoc analysis (**A** and **C**). Scale bar: 50μm.

### Restored Cav1 expression inhibited nitrative stress in *Egln1*^*Tie2Cre*^*/*Cav1^EC-Tg^ mice

To determine whether Cav1 deficiency is responsible for the augmentation of nitrative stress and the pathogenesis of PAH in CKO mice, we generated a mouse model with Cav1 transgenic expression in ECs in the background of CKO mice (*Egln1*^*Tie2Cre*^*/*Cav1^EC-Tg^, *CKO/Tg*)(28) (**Figure 5A**). Western blotting and immunostaining demonstrated that reduced Cav1 expression seen in CKO mouse lungs was restored in *CKO/Tg* mice (**Figure 5B** and **5C**). Restored endothelial Cav1 expression in ECs in CKO mice drastically reduced the levels of ROS/RNS in the lungs of CKO/Tg mice compared to CKO mice (**Figure 6A**). Further examination of NT expression via Western Blotting and immunostaining showed that NT levels were inhibited in the lung and pulmonary vascular ECs of CKO/Tg mice compared to CKO mice (**Figure 6B** and **6C**), suggesting that Cav1 deficiency is responsible for the increased oxidative/nitrative stress in CKO mouse lungs.

**Figure 5.**
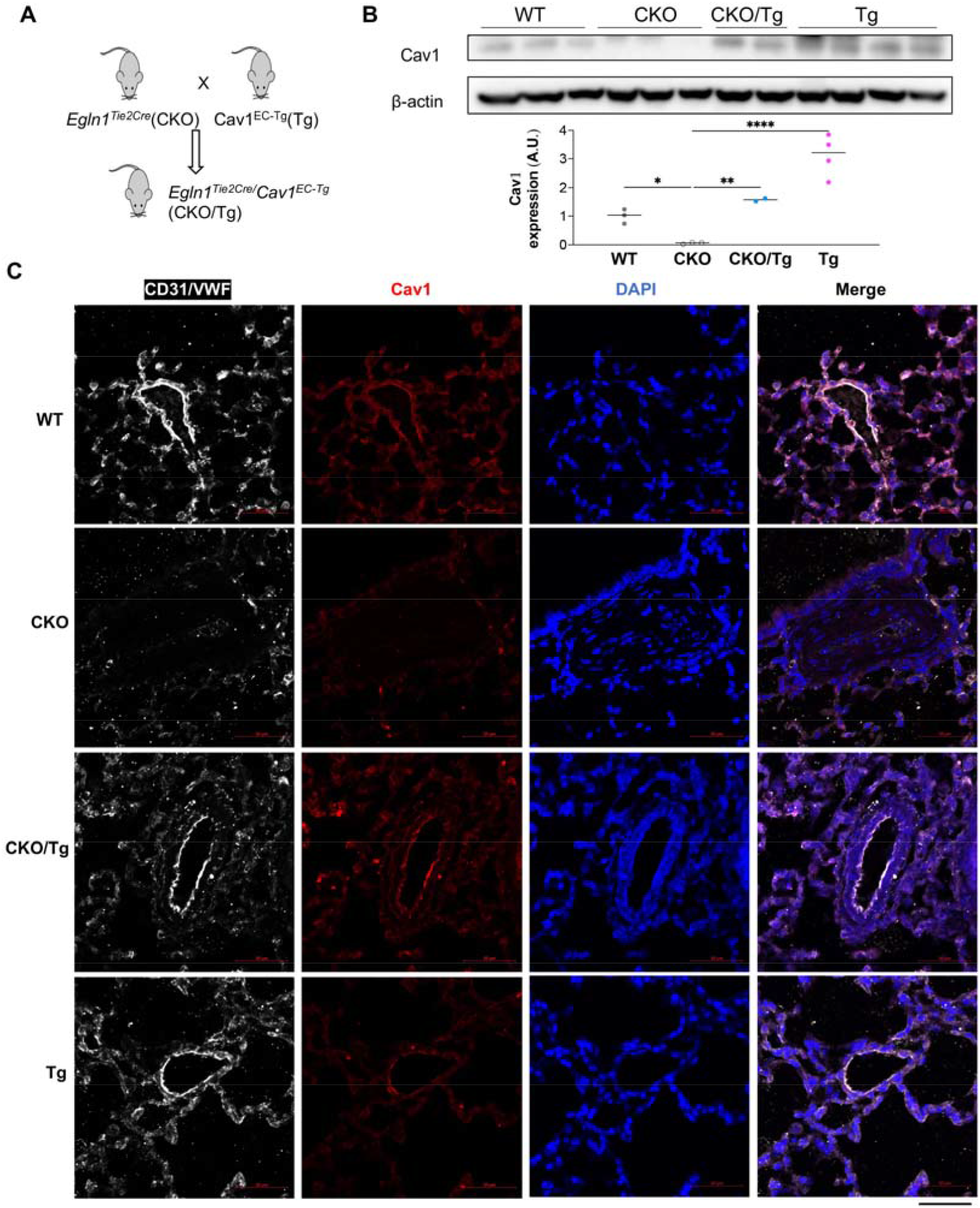
Endothelial Cav1 transgenic expression in *Egln1*^*Tie2Cre*^ mice. (**A**) A diagram showing the strategy of generating *Egln1*^*Tie2Cre*^*/Cav1*^*EC-Tg*^ (CKO/Tg) mice by breeding endothelial Cav1 transgenic mice (*Cav1*^*EC-Tg*^) into the genetic background of CKO mice. (**B**) Western blotting demonstrating restored Cav1 expression in lung from CKO/Tg mice. (C) Immunostaining demonstrated that Cav1 expression was downregulated in lung ECs of CKO mice compared to WT which was restored in CKO/Tg mice. Lung sections were immunostained with anti-Cav1 (red) and CD31/VWF (white). Nuclei were counterstained with DAPI (blue). *, *P* < 0.05, **, *P* <0.01, ****, *P*<0.0001. One-way ANOVA with Tukey post-hoc analysis (**B**).

**Figure 6.**
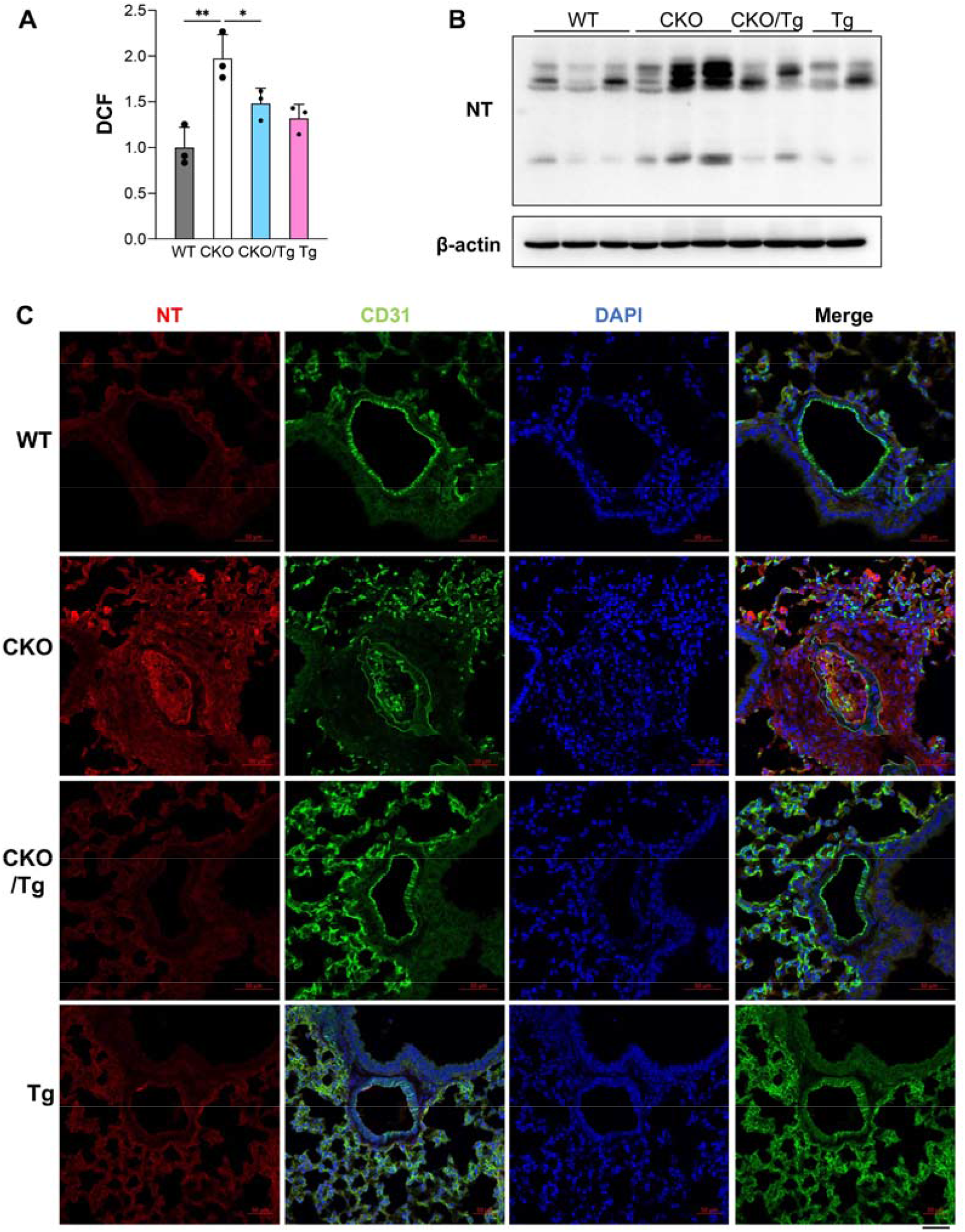
Endothelial Cav1 transgenic expression inhibited excessive ROS/RNS in *Egln1*^*Tie2Cre*^ mouse lungs. (**A**) DCF measurements demonstrating the increased production of ROS/RNS seen in CKO mouse lungs was inhibited in *CKO/Tg* mouse lungs. There is no marked difference in lung generation of ROS/RNS in Tg mice compared to WT mice. WT and CKO data shared with **Figure 1A**. (**B**) Western Blotting demonstrated the reduction of NT levels in *CKO/Tg* mice compared to CKO mice. (**C**) Representative immunostaining showed that CKO/Tg mice exhibited reduced NT levels compared to CKO mice. Lung sections were immunostained with anti-NT (red) and CD31 (Green). Nuclei were counterstained with DAPI (blue). *, *P* < 0.05, **, *P* <0.01. One-way ANOVA with Tukey post-hoc analysis (**A**).

### Cav1 transgenic expression attenuates PAH seen in *Egln1*^*Tie2Cre*^ mice

We next determined if Cav1 transgenic expression in CKO mice will rescue the hypertensive pulmonary phenotype. Hemodynamic measurement demonstrated a partial decrease of RVSP in *CKO/Tg* at age of 3.5 months compared to CKO mice (**Figure 7A**). RV hypertrophy, determined by weight ratio of RV versus left ventricular plus septum (LV+S) was also partially attenuated in *CKO/Tg* mice compared to CKO mice (**Figure 7B**). Pathological examination showed that *CKO/Tg* mice had reduced pulmonary vascular remodeling, including decreases of neointima and occlusive lesions, pulmonary arterial wall thickness, and muscularization of distal pulmonary vessels in comparison of CKO mice (**Figure 7C-7F**).

**Figure 7.**
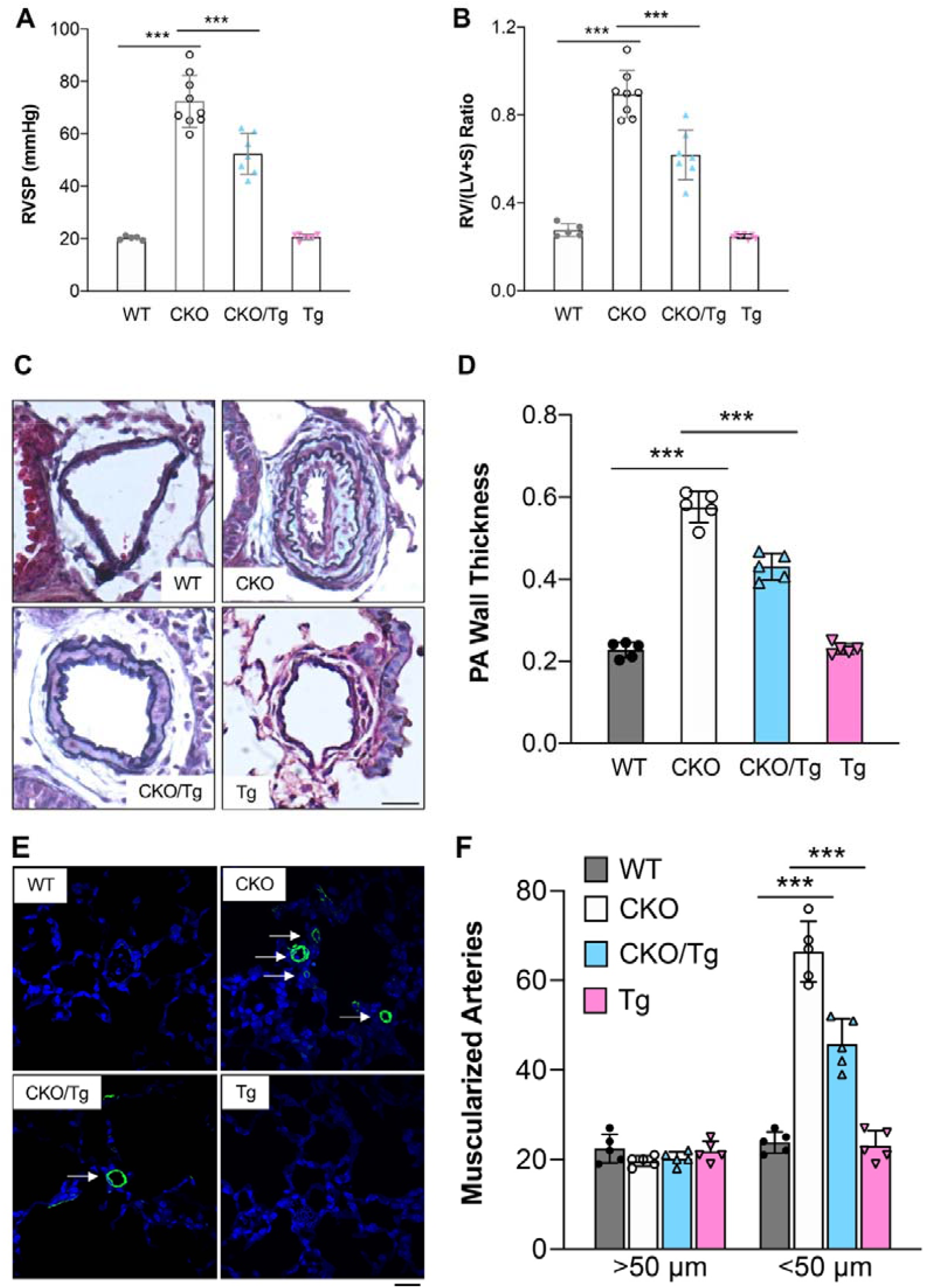
Transgenic expression of Cav1 in ECs inhibited severe PAH seen in *Egln1*^*Tie2Cre*^ mice. (**A, B**) CKO/Tg mice exhibited marked decreases of RVSP (**A**) and RV hypertrophy (**B**) compared to CKO mice at age of 3.5-month old. (**C, D**) Representative micrographs of Pentachrome staining showing inhibited obliterative vascular remodeling including diminished neointima formation and reduced wall thickness in the lung of CKO/Tg mice. (**E, F**) Immunostaining of α-smooth muscle actin (α-SMA) demonstrating inhibition of muscularization of distal pulmonary vessels in the lungs of CKO/Tg mice. Scale bar: 50μm. ***, *P* < 0.001. One-way ANOVA with Tukey post-hoc analysis (**A, B, D** and **F**).

### Cav1 transgenic expression normalizes RV and pulmonary artery functions in *Egln1*^*Tie2*Cre^/Cav1^EC-Tg^ mice

We next evaluate whether Cav1 transgenic expression will inhibit right heart dysfunction seen in CKO mice. As shown in **Figure 8A-8C**, echocardiography showed that *CKO/Tg* mice had marked decreases of RV wall thickness compared to CKO mice. *CKO/Tg* mice also exhibited marked improvements of RV contractility assessed by RV fraction area change (RV FAC) compared to *Egln1*^*Tie2Cre*^ mice (**Figure 8D and Supplemental Video 1-2**). However, we did not observe significant change of heart rate, cardiac output and left ventricular fraction shortening (LV FS) in *CKO/Tg* mice compared with CKO mice (**Supplemental Figure 3A-3C**). Pulmonary artery (PA) function assessed by PA acceleration time/ejection time (AT/ET) showed that CKO/Tg mice exhibited improved PA function compared to CKO mice (**Figure 8E**). These data demonstrated that endothelial Cav1 transgenic expression partially but significantly rescues the defective PA and RV functions seen in CKO mice.

**Figure 8.**
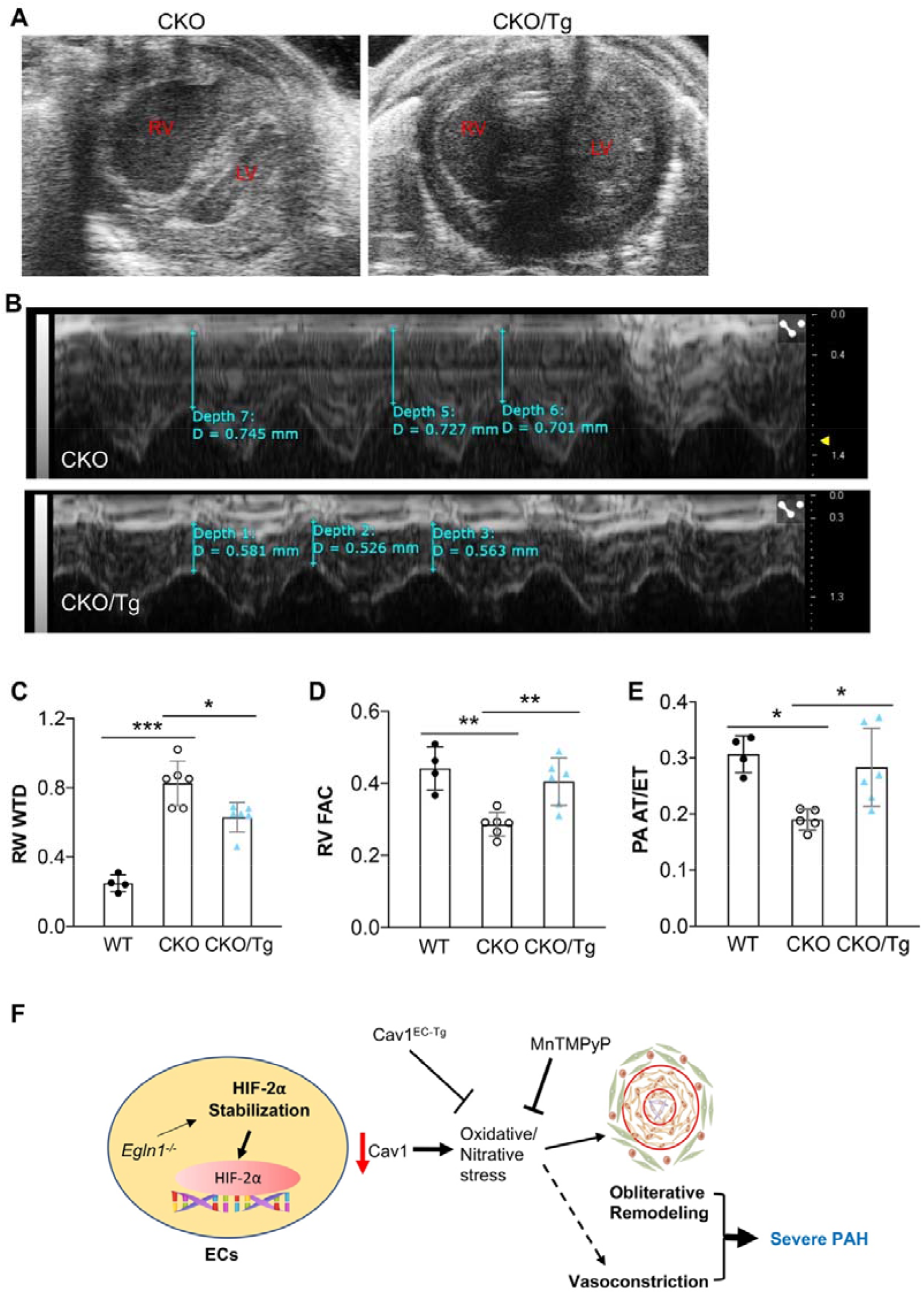
Cav1 transgenic expression in ECs improved right heart and pulmonary artery functions in *Egln1*^*Tie2Cre*^*/*Cav1^EC-Tg^ mice. (**A**) Representative echocardiography B-mode showing reduced RV chamber size in CKO/Tg mice compared to CKO mice. (**B**) Representative echocardiography M-mode showing reduced RV wall thickness at diastolic stage in CKO/Tg mice compared to CKO mice. (**C**) RV wall thickening seen in CKO mice was reduced in CKO/Tg mice. (**D**) RV contractility was improved by restored endothelial Cav1 expression in CKO/Tg mice compared to CKO mice. (**E**) Pulmonary arterial function assessed by PA AT/ET ratio was rescued in CKO/Tg mice. (**F**) A diagram showing that PHD2 deficiency induces decreases of endothelial Cav1 expression resulting in augmented nitrative stress which contributes to obliterative pulmonary vascular remodeling and severe PAH. Restoration of endothelial Cav1 expression or ROS scavenging may be effective therapeutic approaches for the treatment of PAH. *, *P* < 0.05, **, *P* <0.01, ***, *P* < 0.001. One-way ANOVA with Tukey post-hoc analysis (**C**-**E**).

## Discussion

The present study demonstrated that nitrative stress was markedly augmented in the lungs, especially in vascular lesions of CKO mice, recapitulating the pathological feature of clinical PAH. Augmentation of nitrative stress was due to endothelial Cav1 deficiency. Endothelial Cav1 transgenic expression in *CKO/Tg* mice reduced nitrative stress levels and attenuated PAH evident by reduction of RVSP, pulmonary vascular remodeling, and RV hypertrophy, as well as improvements of RV and PA function. We also observed that pharmacological scavenging of ROS/RNS markedly reduced the severity of pulmonary vascular remodeling and PAH in CKO mice. These findings suggest that restoration of Cav1 expression in ECs and ROS/RNS scavenging represent therapeutic strategies for the treatment of severe PAH, including IPAH (**Figure 8F**). The clinical relevance is validated by our previous observation that PHD2 expression was diminished in ECs of occlusive pulmonary vessels in IPAH patients(25).

Prominent oxidative/nitrative stress is a pathological feature of clinical PAH (12, 17). Our recent study has for the first time shown that CKO mice develop spontaneously severe PAH with obliterative pulmonary vascular remodeling including occlusive vascular lesions and formation of plexiform-like lesions and marked elevation of RVSP ranging from 70-100 mmHg as well as development of right heart failure (25). Here we show a marked increase of ROS/RNS in CKO mouse lungs and prominent anti-NT staining indicative of nitrative stress in pulmonary vascular lesions, especially in ECs as well as inflammatory cells. This study provides further evidence of CKO mice as an excellent mouse model of human PAH. Furthermore, treatment of the mice with the ROS scavenger Mn inhibited occlusive pulmonary vascular remodeling and attenuated PAH, demonstrating the pathogenic role of oxidative/nitrative stress in contributing to the severity of PAH.

Nitrative stress is formed by excessive superoxide and NO which chemically react to form peroxynitrite. To specifically inhibit peroxynitrite formation, we also tried to reduce NO production by treating the CKO mice with a NO synthase inhibitor L-NAME (1 mg/ml in drinking water). However, all the 5 CKO mice died within ∼7 days treatment, suggesting NO is essential for the maintenance of vascular homeostasis and survival of these mutant mice. It is possible that the prominent nitrative stress seen in pulmonary vascular lesions greatly depletes NO leading to limited NO bioavailability in CKO mice, thus, NOS inhibition further lowers NO bioavailability which causes high mortality.

Previous studies have shown decreases of *CAV1* mRNA and protein levels in lung ECs of patients with PAH(20, 30, 31). Cav1 deficiency is also evident in the arterial lesions of PAH rats including both Sugen5416/hypoxia-induced PAH and monocrotaline-induced PAH(34, 35). Our previous study provides the first genetic evidence that Cav1 deficiency induces PH in mice (19). Others also observe pulmonary defects in *Cav1*^*-/-*^ mice (36). A recent study shows that Cav1 deficiency in ECs promote more severe PH in response to hypoxia (37). Consistently, *CAV1* mutation is associated with heritable PAH in patients (38). Together, these studies provide unequivocal evidence of CAV1 deficiency in the pathogenesis of PAH. Our other study has shown that mice with genetic deletions of both *Cav1* and *Nos3* or treatment of *Cav1*^-/-^ mice with either NO Synthase inhibitor L-NAME or ROS/RNS scavenger Mn inhibited PAH, suggesting chronic activation of eNOS secondary to Cav1 deficiency is the mechanism of PAH induced by CAV1 deficiency (21). Others also show that L-NAME treatment inhibits PH in *Cav1*^*-/-*^ mice (39). eNOS activation-derived NO reacts with superoxide to form the damaging peroxynitrite which causes posttranslational modification of proteins such as PKG via tyrosine nitration in *Cav1*^-/-^ mice. PKG nitration impairs its kinase activity and thereby induces pulmonary vascular remodeling and PH (21, 23). Accordingly, *Prkg1*^*-/-*^ (i.e., PKG deficiency) mice also exhibit PH (23). PKG nitration is prominent in lung tissues of IPAH patients (21, 24). These studies provide unequivocal evidence that Cav1 deficiency induces nitrative stress leading to PKG nitration and thus PH.

In CKO mice, we observed elevated levels of ROS/RNS, PKG nitration and decrease of endothelial Cav1 expression. We demonstrated that Cav1 deficiency in CKO mice is involved in the regulation of nitrative stress level as endothelial Cav1 transgenic expression normalizes the levels of ROS/RNS and inhibits nitrative stress evident by diminished NT levels in pulmonary vascular ECs and thus inhibits obliterative pulmonary vascular remodeling and reduces PAH as well as improves RV and PA functions in *CKO* mice. Especially, we observed complete inhibition of occlusive vascular remodeling without neointima formation in both CKO/Tg mice and Mn-treated CKO mice. The medial thickness was less affected in these mice, which is consistent with the markedly reduced but still elevated RVSP. Together, our data demonstrate that endothelial Cav1 deficiency secondary to PHD2 deficiency mediates the augmentation of nitrative stress in CKO mouse lungs and pulmonary vascular lesions and contributes to the severity of PAH.

Given the limitation that both *CKO* mice and Cav1^EC-Tg^ mice are not conditional, thus the effects of these transgenic manipulations would be exerted throughout developmental stage. It is possible that the phenotype may reflect effects on embryonic vascular development. Since PAH is a disease that occurs mainly in adults (and less commonly in children), future studies are warranted to address these mechanisms using similar manipulations postnatally.

In summary, we have demonstrated prominent nitrative stress in vascular lesions of CKO mice, which is mediated by endothelial Cav1 deficiency ascribed to HIF-2α activation. ROS/RNS scavenger treatment or transgenic expression of endothelial Cav1 inhibits obliterative pulmonary vascular remodeling and attenuates PAH in CKO mice. Thus, Cav1 deficiency-induced oxidative/nitrative stress secondary to PHD2 deficiency is a part of the mechanisms of severe PAH in CKO mice as seen in PAH patients.

## Acknowledgement

We greatly appreciate Dr. William C. Sessa from the Departments of Pharmacology, and Medicine at Yale School of Medicine for his generosity of providing us the *Cav1*^*EC-Tg*^ mice.

## Author contributions

Z.D. and Y.Y.Z. conceived the experiments. Z.D., J.D., B.L., D. Y., Y. P. and M.M.Z designed, carried out experiments, and analyzed the data. Z.D., R.V. and Y.Y.Z. analyzed and interpreted the data. Z.D. wrote the manuscript. Z.D. and Y.Y.Z. supervised the project and revised the manuscript. All authors approved the manuscript.

## Supplemental Figure legends

**Supplemental Figure 1.**
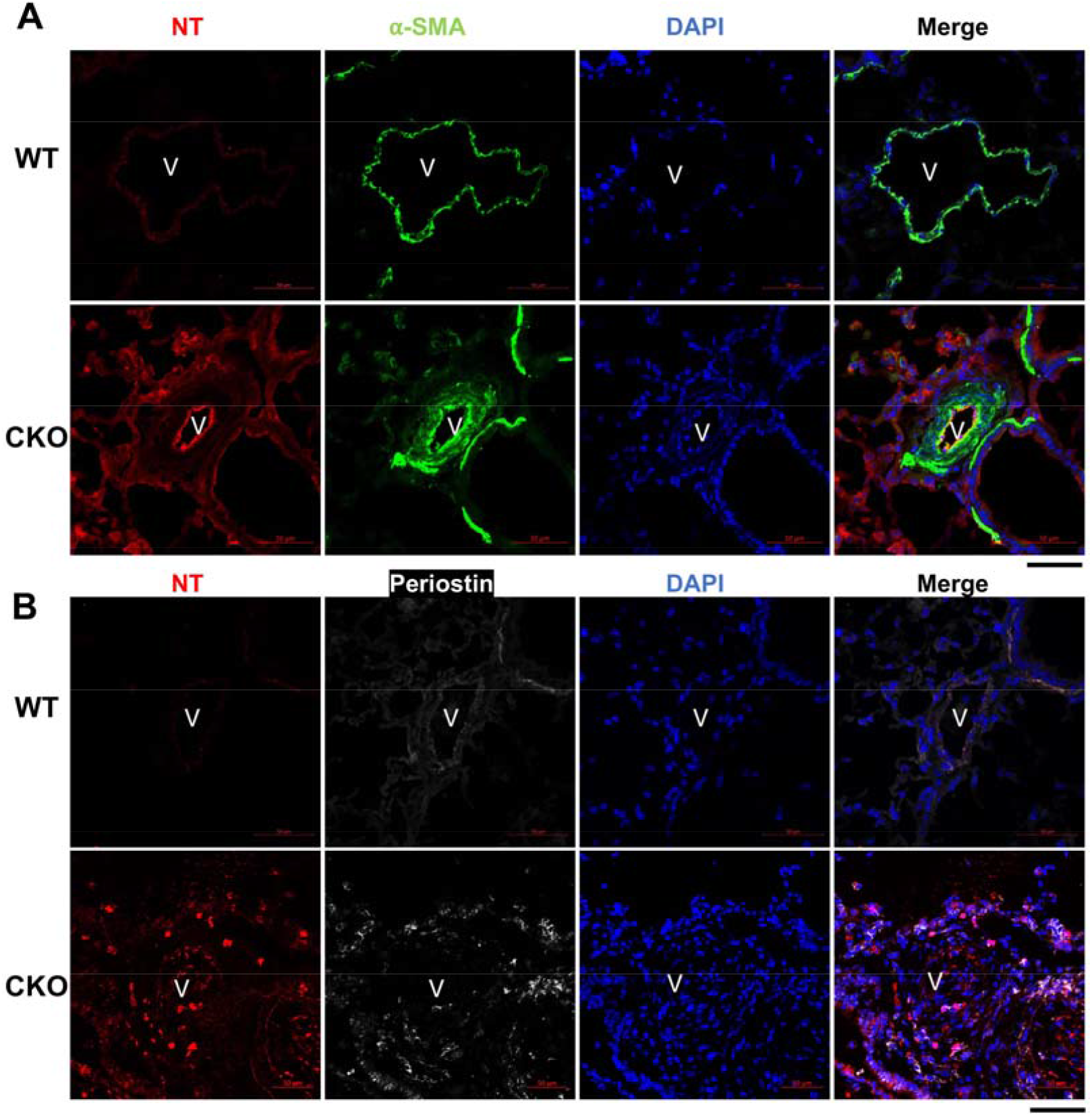
Nitrative stress in smooth muscle cells and fibroblasts in the pulmonary vascular lesions of *Egln1*^*Tie2Cre*^ mice. Representative micrographs of immunostaining showing relatively low levels of NT expression in smooth muscle cells and fibroblasts in pulmonary vascular lesions of CKO mice (3.5 months old). Lung sections were immunostained with anti-NT antibody (red) for detection of nitrative stress. Anti-α-SMA antibody was used to label smooth muscle cells (green, **A**). Anti-Periostin antibody was used to label fibroblasts (white, **B**). Nuclei were counterstained with DAPI (blue). The strong positive NT signal in (A) was from pulmonary vascular ECs not SMCs.

**Supplemental Figure 2.**
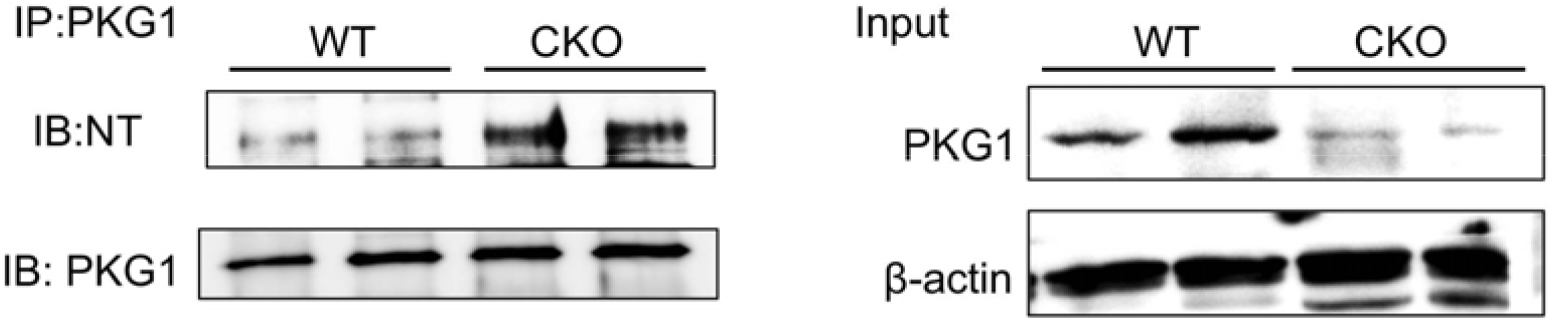
Immunoprecipitation assay demonstrating that PKG nitrotyrosine modification was markedly increased in *Egln1*^*Tie2Cre*^ mice. Lung tissues were collected from 3.5 months old mice for homogenization. 450 μg of lysates/mouse were then immunoprecipitated with anti-PKG antibody and blotted with anti-NT antibody. The same membrane was also blotted with anti-PKG antibody as a loading control.

**Supplemental Figure 3.**
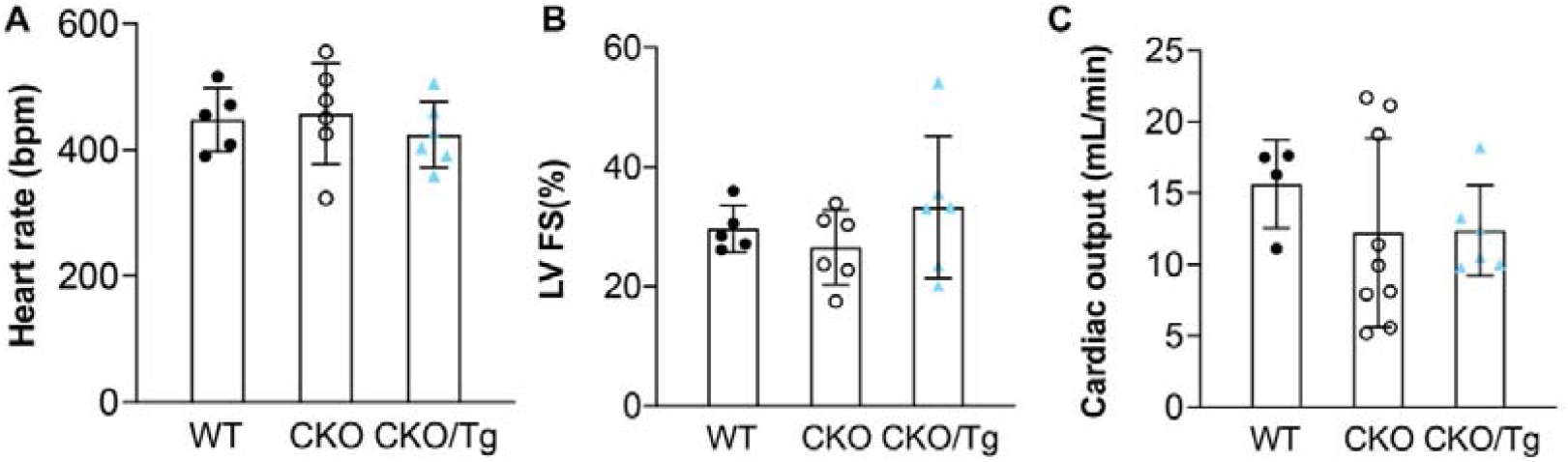
Echocardiography analysis showing similar left heart function and cardiac output in CKO/Tg mice and CKO mice. At age of 3.5 months, the mice were subject to echocardiography to assess heart rate (**A**), left ventricular fraction shortening (LV FS), indicative of contractility (**B**), and cardiac output (**C**). bpm= beats per minute. One-way ANOVA with Tukey post-hoc analysis (**A**-**C**).

**Supplemental Video 1, Echocardiography M-Mode of CKO mice**.

DOI: 10.6084/m9.figshare.16674463

**Supplemental Video 2, Echocardiography M-Mode of *CKO/Tg* mice**.

DOI: 10.6084/m9.figshare.16674478

